# Topological embedding and directional feature importance in ensemble classifiers for multi-class classification

**DOI:** 10.1101/2024.08.01.605982

**Authors:** Eloisa Rocha Liedl, Shabeer Mohamed Yassin, Melpomeni Kasapi, Joram M. Posma

## Abstract

Cancer is the second leading cause of disease-related death worldwide, and machine learning-based identification of novel biomarkers is crucial for improving early detection and treatment of various cancers. A key challenge in applying machine learning to high-dimensional data is deriving important features in an interpretable manner to provide meaningful insights into the underlying biological mechanisms

We developed a class-based directional feature importance (CLIFI) metric for decision tree methods and demonstrated its use for the The Cancer Genome Atlas proteomics data. We incorporated this metric into four algorithms, Random Forest (RF), LAtent VAriable Stochastic Ensemble of Trees (LAVASET), and Gradient Boosted Decision Trees (GBDTs), and a new extension incorporating the LAVA step into GBDTs (LAVABOOST). Both LAVA methods incorporate topological information from protein interactions into the decision function.

The different models’ performance in classifying 28 cancers resulted in F1-scores of 93% (RF), 92% (LAVASET), 89% (LAVABOOST) and 86% (GBDT), with no method outperforming all others for individual cancer type prediction. The CLIFI metric allowed the visualisation of the model decision making functions, and the distributions indicated heterogeneity in several proteins (MYH11, ER*α*, BCL2) for different cancer types (including brain glioma, breast, kidney, thyroid and prostate cancer).

We have developed an integrated, directional feature importance metric for multi-class decision tree-based classification models that facilitates interpretable feature importance assessment. The CLIFI metric can be used in conjunction with incorporating topological information into the decision functions of models to add inductive bias for improved interpretability.

**Availability:** All codes are available for data curation from https://github.com/EloisaRL/TCGA-proteomics-pipeline and the LAVASET (v1.0) package from https://github.com/melkasapi/LAVASET.

## Introduction

Cancer has risen from third place in 2010 to second place in 2019 as the leading cause of disease-related death worldwide, and it is forecasted that the global cancer burden will continue to grow for the next two decades (1–3). As a result, the application of machine learning (ML) methods in oncology has expanded (4). In addition to data analysis automation, ML models often show higher accuracy in diagnosis and survival predictions than traditional clinical methodology (5, 6). Given the shift in cancer medical practice to personalised and targeted treatments, classification of cancer patients is moving towards the need of identifying subpopulations within one cancer type (7, 8).

Ensemble methods, a subfield of ML, are commonly used for this classification task (9–11), as they excel in maintaining performance with high-dimensional datasets that have a small number of samples relative to the number of features (small *n*, large *p*), unlike other ML methods - including deep learning algorithms that require more samples for effective training (12). Such high-dimensional datasets are common in cancer research, including genomic, epigenetic, transcriptomic, proteomic, and clinical data. Examples of ensemble methods include Random Forests (RFs) (13) and Gradient Boosted Decision Trees (GBDTs) (14). Despite gradient boosting showing superior prediction performance in previous cancer prediction studies compared to RF, support vector machines, and logistic regression algorithms (15–17), the existence of heterogeneity limits the application of RF and GBDT in cancer. While binary classification models can still be easily interpreted in how predictions are being made, by using Gini values, multi-class classification algorithms are not (18).

High-dimensional datasets often contain correlated features (feature interactions), which can lead the model to overfitting, by capturing noise or redundant features. Hence, they compromise model generalisation by assigning importance to non-biologically meaningful features (18). There are techniques that can address the problem of correlated features e.g. Boruta algorithm (19), a novel RF algorithm, but they do not explicitly incorporate domain knowledge of correlations. The Boruta algorithm was utilised to identify microRNAs distinguishing normal and ovarian cancer patients, yielding more differentially expressed microRNAs compared to previous studies (20). Subsequent pathway analysis validated the ML findings, demonstrating concordance with existing literature (20). While these algorithms are effective, biological pathway interpretation and validation are necessary to verify and interpret the results. Topological ML algorithms incorporating this type of information into the model architecture can address the multicollinearity problem and make the model more interpretable thereby reducing the need for extensive post hoc validation analyses.

Informative feature importance values depend on two factors: class-specificity and directionality. ML models in cancer research commonly utilise binary classification models with Gini-based feature importance assignment, which considers only the magnitude of features’ influence on predictions (10, 21, 22). Applying multi-class classification algorithms offer greater versatility as they enable prediction of multiple classes using a single model.

In cancer research, ensemble methods with RF and GBDTs are rarely used for multi-class classification, instead a binary one versus all approach is usually employed to identify class-specific important features. For example, Ortiz-Ramon et al. used two models to discriminate brain metastases patients, based on their primary site of origin: a multiclass RF model with 87% accuracy and a one versus all model with 82% accuracy (23). They then, identified class-specific Gini-based feature importance by analysing the one vs all models. While the Gini-based feature importance did help identify differentiating biomarkers, it did not clarify whether high or low values of the biomarkers are associated with each class. The need for class-specific and directional feature importance assignment is essential for broader application of multi-class ensemble models. Directional feature importance calculations are available for ensemble models, for example permutation importance (13) and SHapley Additive exPlanations (SHAP) (24); however, they are post-hoc and not incorporated within the methods. To the best of our knowledge, no such method exists for RF or GBDT that is integrated within algorithm.

Our contribution aims to address two gaps. The first is to introduce an integrated, directional feature importance metric for decision tree-based models (such as RF and GBDT) to facilitate feature importance assessment for multi-class classification. We demonstrate the feature importance measure on the small-scale Fisher Iris data before applying it on a large dataset. The second is to expand on recently published work, LAVASET (LAtent VAriable Stochastic Ensemble of Trees) (25), that demonstrated how data-specific properties can be integrated in the ensemble model for better interpretability for correlated, temporal and spatial features. We extend this paradigm to GBDTs, producing LAVABOOST (LAtent VAriable gradient BOOSTed decision trees), and apply the 4 algorithms for multi-class classification in The Cancer Genome Atlas (TCGA) proteomics dataset (26) of 28 different cancers with incorporation of topological information of protein-protein interactions.

## Materials and Methods

### Data

#### Data - Fisher Iris

The original Fisher Iris data was obtained from `scikit-learn` and used to demonstrate the directional feature importance. Here, we added 3 random noise features of different distributions (Gaussian, uniform, and bimodal) to the original features (sepal length, sepal width, petal length, petal width) to test the feature importance for features with different conditions of (random) distributions (we refer to this dataset as ‘Iris’). Additionally, we randomly shuffled the values for each feature to break the correlation pattern of the Iris data to demonstrate the directional feature importance output for models without predictive power, this dataset is referred to as ‘Iris-permuted’.

#### Data - TCGA

Proteomic profiling data of 28 cancer types was obtained from TCGA project. These publicly available datasets were generated from tumour tissue analysis conducted by the University of Texas MD Anderson Cancer Centre using reversed-phase protein arrays (26). A total of 243 glioblastoma multiforme (GBM), 435 brain lower grade glioma (LGG), 354 head and neck squamous cell carcinoma (HNSC), 381 thyroid carcinoma (THCA), 126 oesophageal carcinoma (ESCA), 226 sarcoma (SARC), 352 skin cutaneous melanoma (SKCM), 365 lung adenocarcinoma (LUAD), 328 lung squamous cell carcinoma (LUSC), 62 mesothelioma (MESO), 90 thymoma (THYM), 919 breast invasive carcinoma (BRCA), 357 stomach adenocarcinoma (STAD), 184 liver hepatocellular carcinoma (LIHC), 120 pancreatic adenocarcinoma (PAAD), 46 adrenocortical carcinoma (ACC), 82 pheochromocytoma and paraganglioma (PCPG), 478 kidney renal clear cell carcinoma (KIRC), 63 kidney chromophobe (KICH), 216 kidney renal papillary cell carcinoma (KIRP), 363 colon adenocarcinoma (COAD), 132 rectum adenocarcinoma (READ), 432 ovarian serous cystadenocarcinoma (OVCA), 440 uterine corpus endometrial carcinoma (UCEC), 172 cervical squamous cell carcinoma and endocervical adenocarcinoma (CESC), 352 prostate adenocarcinoma (PRAD), 122 testicular germ cell tumours (TGCT), 343 bladder urothelial carcinoma (BLCA), were included in the initial dataset. All samples were obtained from the TCGA portal (version 39.0, release date 04/Dec/2023) at https://portal.gdc.cancer.gov/.

#### Data Processing - TCGA

Proteins with more than 50% missing values were removed. Missing values were imputed using the k-Nearest Neighbours Imputer (`scikit-learn, v1.3.2`) with k=5. The protein list was further refined to include only those that had been validated by Western blotting in a TCGA pan-cancer independent study (27). Compared to the prior study, we included additional cancer types. To assess whether this affected clustering ability, an unsupervised analysis of the data with and without the additional cancer types showed no clear differences. Consequently, the final dataset comprised of 7,783 samples with 113 proteomic features.

### Machine Learning Classifiers

#### Random Forest and LAVASET

The LAVASET package provides implementation for both the RF and LAVASET algorithms (25), based on the principles of Breiman’s original RF algorithm (13). LAVASET utilises the original `C++` implementation for computational efficiency, with main functions written in `Python` for easier user readability. The traditional Classification And Regression Trees (CART) algorithm (28) is used to build individual decision trees. The modifiable hyperparameters for both algorithms are the number of trees, number samples per tree, and the maximum number of features. LAVASET in addition has a distance parameter which defines the threshold strength of feature interactions to be considered in the embedding.

The ‘LAVA’ step occurs before the feature is chosen for the best split (see (25) for complete details). In brief, Principal Component Analysis (PCA) is used to identify the first right singular vector (loading) of the decomposition of the feature selected and its correlated features. The loading is then used to transform the original features into a single score (left singular value). This is performed for all features (and their neighbours/correlated features) selected at the splitting step. Therefore, the feature dataset is transformed to consist of latent variables instead of the original feature values.

#### Extension of Latent Variable Embedding to Boosting Algorithms

The multiclass classification gradient boosting decision tree (GBDT) algorithm developed in this study was adapted from Matt Bowers’ implementation of Algorithm 6 from greedy function approximation (14). This implementation uses a one-against-all approach to reduce the problem into K binary problems, where K is the number of classes. Our implementation of GBDT, as well as LAVABOOST (Latent Variable embedding into GBDTs), uses the traditional CART algorithm (28) with the original `C++` implementation to build individual decision trees rather than the `scikit-learn DecisionTreeRegressor` function. The sequential trees in GBDTs and LAVABOOST predict the errors of the previous prediction therefore, to prevent biases, the predictions are initialised to 0.

LAVABOOST has a number of prerequisites and hyperparameters that can be optimised (see section). A distance matrix is provided by the user containing the strength of interactions between features in the input data set (see section). The features the algorithm considers correlated are those with an interaction strength less than or equal to the threshold set by the user, the threshold is set by the ‘distance’ parameter. The modifiable optimisation parameters include the number of estimators (boosting rounds), learning rate, number samples per tree, and the maximum number of features. When the number of samples per tree or maximum number of features is set to `None`, all samples/features, respectively, are used.

#### Distance Metric for Latent Variable Embedding

For use in LAVASET and LAVABOOST only, we produced a distance matrix representing the protein interactions. The Search

Tool for the Retrieval of Interacting Genes/Proteins database (STRINGdb) version 12 (https://string-db.org/) (29) was used to identify the protein-protein interactions of all proteins in the TCGA dataset. For any TCGA protein name not found in the STRINGdb database, we used the gene name of each protein (listed in (27)) to find protein interactions between all 113 features. Protein interactions were defined by self-interactions (distance d=0), proteins linked to the same gene (d=1), protein-coding gene-gene interactions (d=2, evaluated using confidence level of 0.7), and no interaction (d=100).

#### Directional Feature Importance

The directional features importance calculation occurs after the optimal feature (or latent feature for LAVASET and LAVABOOST) has been selected with the optimal split. For all splits, the Gini coefficient values are calculated. For RF and GBDT, the Gini is attributed to the chosen feature for the optimal split. For LAVASET and LAVABOOST, a normalised Gini feature importance is calculated for each original feature using the absolute value of the loadings (normalised to a sum of 1). Below we describe the newly proposed CLass-based Integrated directional Feature Importance (CLIFI) calculation for ensemble models such as RF, GBDT, LAVASET and its novel extension LAVABOOST. The benefit of this calculation is that it is integrated into the algorithm and therefore feature importances of the models can be directly inferred without need for additional (post-hoc) techniques.

The G-test (Eqn. 1), a complimentary approach to Kullback-Leibler divergence, to compare the observed and expected frequencies of categorical data (i.e. the distribution of groups after splitting). Where *O*_*i*_ is the observed count for the i^th^ category and likewise, *E*_*i*_ is the expected count for the same category under the null hypothesis.

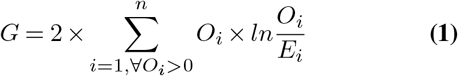

We use a modified version of the G-test to quantify the distribution of samples in a class in each of the splits, where the observed count in Equation 1 is replaced with the frequency in the left split (*L*_*i*_) for class i (Eqn. 2), and the expected count is replaced by half the frequency in the parent node 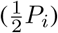. The same process is done for frequency in the right split (*R*_*i*_, Eqn. 3). These two quantities are partial G-tests (*M*_*ij*_), and summing these results is the combined G-test value for that class. This assumes that *P*_*i*_, and either (or both) of *L*_*i*_ and *R*_*i*_, are non-zero.

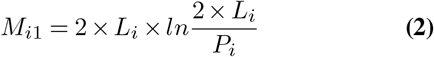

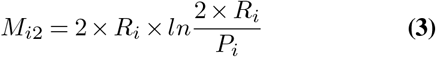

A perfect split is represented by 2 *× P*_*i*_ *× ln*2, therefore we normalise *M* by the perfect split to scale it between 0 (equal split) and 1 (perfect split) and subsequently multiply with *P*_*i*_*/A*_*i*_ to account for the number of samples of the class considered for the split relative to all samples in that tree (*A*_*i*_, ‘ancestor’ node). The directionality of the feature for class *i* is indicated by the sign of *M*_*i*2_ − *M*_*i*1_. Combining this, and simplifying, results in Equation 4 for the CLIFI of feature *j* for class *i*:

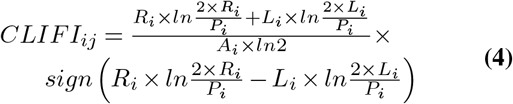

A positive CLIFI value for a feature signifies that the direction of the split is weighted more towards the right node (higher split values) than the left (lower split values). The magnitude of the association is given by the numerical value itself, where the closer is it to 1 (or −1), the more it is associated with higher (or lower) values of that feature for RF and LAVASET models. The aggregated CLIFI (aCLIFI) value is the sum of the CLIFI values across all trees for a specific feature and class, and for comparison between models the normalised aggregated CLIFI (naCLIFI) values represent a division by the highest absolute aggregated CLIFI value across all features and classes.

For GBDT and LAVABOOST the above cannot be used directly because each class label (*i*) predicted by the model is an error label, therefore, samples of the same class (noted by *h*) may have different error labels. Therefore, the CLIFI for these models is calculated by scaling the CLIFI value for the error label by the proportion of samples in the real class (Eqn. 5), where *x* represents the distinct error labels associated with class *h*, and *S* is the number of samples within class *h* with error label *i* divided by the total number of samples in class *h*.

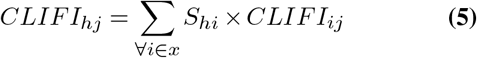

#### Model Evaluation

30% of the data was set aside for testing, with the remaining 70% split 80:20 into training and validation sets. Individual class frequencies were balanced across the subsets of data. The training set (ca. 56% of the total data) was used to calculate individual models with different hyperparameter settings, and the validation set (ca. 14% of the total data) was used to determine the optimal hyperparameters. The optimal model was then used to predict the left-out test set (30%). For the TCGA data, RF, LAVASET, and GBDT were run 100 distinct times and LAVABOOST 10 distinct times with different random states to evaluate the model robustness.

The optimal hyperparameters for RF and LAVASET models were models with 150 trees and ‘sqrt’ of number of features for each split. The values trialed for number of trees were 100 to 180 in steps of 10. Both models used 80% of samples per tree. Optimal parameters for GBDT and LAVABOOST were 130 estimators (boosting rounds), a learning rate of 0.1, ‘sqrt’ of number of features for each split, and including all samples (‘None’). The values trialed for number of trees were 10 to 150 in steps of 10, and the values trialed for learning rate were 0.1, 0.2, and 0.3. The distance parameter for LAVABOOST and LAVASET was set to 2.

Classification performance was assessed based on accuracy, precision, recall and F1-score, with the mean ± standard deviation reported for each model, for the test set. The proximity matrix was used as input to UMAP (number of components=`2`, init=`random`, random state=`0`) (30) to visualise the similarity between samples from different cancer types. The Iris dataset was split 80:20 into training and validation sets. The optimal parameters for the Iris data were ‘sqrt’ of number of features per split for RF and GBDT, 100 trees for RF, and 7 boosting rounds with 0.1 learning rate for GBDT.

#### Evaluation and Visualisation of CLIFI Values

Class feature importance assignment was evaluated for LAVASET, RF, LAVABOOST, and GBDT using the python package NetworkX. A template protein layout on a hexagonal grid was defined based on the protein interactions, thus interacting proteins are closer together, and was used for all algorithms. The feature importance results present the top 10 proteins with the highest CLIFI value (and higher than the average CLIFI value of the positively assigned proteins), with the same was done for the negatively assigned proteins (10 lowest CLIFI values that are lower than the average CLIFI of negative CLIFI values), with LAVASET and LAVABOOST also showing protein interactions between these proteins. For visualisation purposes that was restricted to the top 10 to allow all edges to be drawn without node-edge crossings.

Paired and unpaired t-tests were used to assess differences between methods in model performance. A Kruskal-Wallis test was used to evaluate whether features’ CLIFI values were significantly different between classes. For those features with significant differences, a Mann-Whitney U test (Wilcoxon rank sum test) was used for pairwise comparisons and these were visualised in a heatmap. All p-values were adjusted for multiple testing using the Benjamini-Hochberg method.

#### Compute, Software Versioning, and Code Availability

All calculations were performed on an HP Z6 G4 workstation with 16-core Intel(R) Xeon(R) Silver 4110 CPU @ 2.10GHz with 128GB RAM.

Combining the TCGA study datasets and defining the protein interactions were implemented with R in (v4.3.2) R Studio (v2023.09.1+494). These codes are available from https://github.com/EloisaRL/TCGA-proteomics-pipeline.

All other analyses were implemented in Python (v3.12.1) with libraries and packages: `lavaset` (v0.1.1), `matplotlib` (v3.8.2), `networkx` (v3.2.1), `numpy` (v1.26.2), `pandas` (v2.1.4), `plotly` (v5.19.0), `scikit-learn` (v1.3.2), `seaborn` (v0.13.2), `umap` (v0.1.1), and `umap-learn` (v0.5.5).

The code for GBDT and LAVABOOST, and implementation of the direction feature importance for all 4 algorithms (including RF and LAVASET), have been added as separate branch to the `LAVASET` repository (https://github.com/melkasapi/LAVASET, v1.0.0).

## Results

### Classification Performance

#### Iris Data

We show that our algorithms show consistent performance with respect to the example datasets, where the permuted version has low classification r ates, a nd f or t he real data we show that RF outperforms GBDT (p < 0.0001) (Table 1). For the demonstration of the CLIFI values, we choose to show these for the RF models for both the Iris (predictive) and Iris-permuted (not predictive) datasets.

**Table 1.** Classification performance of RF and GBDT for the Iris and Iris-permuted datasets. Scores for accuracy, precision (weighted), recall (weighted), F1-score (weighted) are given across 20 random initialisations of the models. The RF was run with 100 trees, GBDT with 7 boosting rounds and learning rate of 0.1. Values shown are the mean ± standard deviation, represented as percentages. Values in bold indicate the highest performances for each metric.

| Dataset | Algorithm | Accuracy (%) | Precision (%) | Recall (%) | Macro F1 (%) |
| --- | --- | --- | --- | --- | --- |
| Iris | RF | **90.00** ± 0.00 | **92.73** ± 0.00 | **90.00** ± 0.00 | **91.34** ± 0.00 |
| Iris-permuted | RF | 38.50 ± 2.68 | 50.65 ± 7.00 | 38.50 ± 2.68 | 43.68 ± 4.25 |
| Iris | GBDT | 72.33 ± 1.53 | 72.39 ± 1.12 | 72.33 ± 1.53 | 72.36 ± 1.27 |
| Iris-permuted | GBDT | 29.00 ± 2.13 | 26.80 ± 4.23 | 29.00 ± 2.13 | 27.77 ± 3.26 |

#### TCGA Data

For the TCGA dataset (28 classes), RF outperformed the other methods (Table 2) though while its difference with LAVASET is significant (p < 0 .001) t he difference is small. Both RF and LAVASET outperform GBDT and LAVABOOST, with LAVABOOST outperforming GBDTs (p < 0.001). LAVABOOST appears to have larger variability in its performance compared to LAVASET. The class-specific performance varies between 0% to 100% for all algorithms (see), with no model consistently outperforming other models on all cancer types. For example, while RF and LAVASET have a considerable higher overall accuracy, for glioblastoma, brain lower grade glioma, thyroid, and testicular germ cell cancer the boosting methods have better performance.

**Table 2.** Classification performance of RF, LAVASET, GBDT, and LAVABOOST. Scores for accuracy, precision (weighted), recall (weighted), F1-score (weighted) across 100 random initialisations of RF, LAVASET, and GBDT, and 10 random initialisations of LAVABOOST. Values shown are the mean ± standard deviation. Values in bold indicate the highest performances for each metric.

| Algorithm | Accuracy (%) | Precision (%) | Recall (%) | Macro F1 (%) | Time/model (min) |
| --- | --- | --- | --- | --- | --- |
| RF | **93.07** ± 0.11 | **92.20** ± 0.35 | **93.07** ± 0.11 | **92.63** ± 0.22 | 3.15 |
| LAVASET | 92.35 ± 0.14 | 91.76 ± 0.39 | 92.35 ± 0.14 | 92.05 ± 0.26 | 2.59 |
| GBDT | 86.52 ± 0.22 | 86.33 ± 0.26 | 86.52 ± 0.22 | 85.65 ± 0.26 | 33.58 |
| LAVABOOST | 89.31 ± 0.18 | 89.36 ± 0.48 | 89.31 ± 0.18 | 89.33 ± 0.32 | 216.57 |

### Feature Importance Assessment

#### Iris Data

The CLIFI value importance assignment was first tested on the Iris dataset to serve as a benchmark for demonstrating the interpretability. Figure 1A shows the output from using the Gini coefficient indicating t hat sepal length (SL), petal length (PL), and petal width (PW) are most predictive. In the CLIFI distribution plot (Fig 1B) PL, PW and SL have the broadest spread of CLIFI values, i.e. they deviate most from 0, hence these are likely to be predictive in the model. Figure 1C-E shows the distribution of CLIFI values for the 3 noise variables for each class (Setosa, Versicolour, Virginica) is centered at 0, and while come noise variables had higher Gini coefficients than the real sepal width (SW) feature there are no significant differences between groups in the CLIFIs. The largest differences are seen for the real variables (Fig 1F-I). These distributions clearly show the relationship between individual features and the class information.

**Fig. 1.**
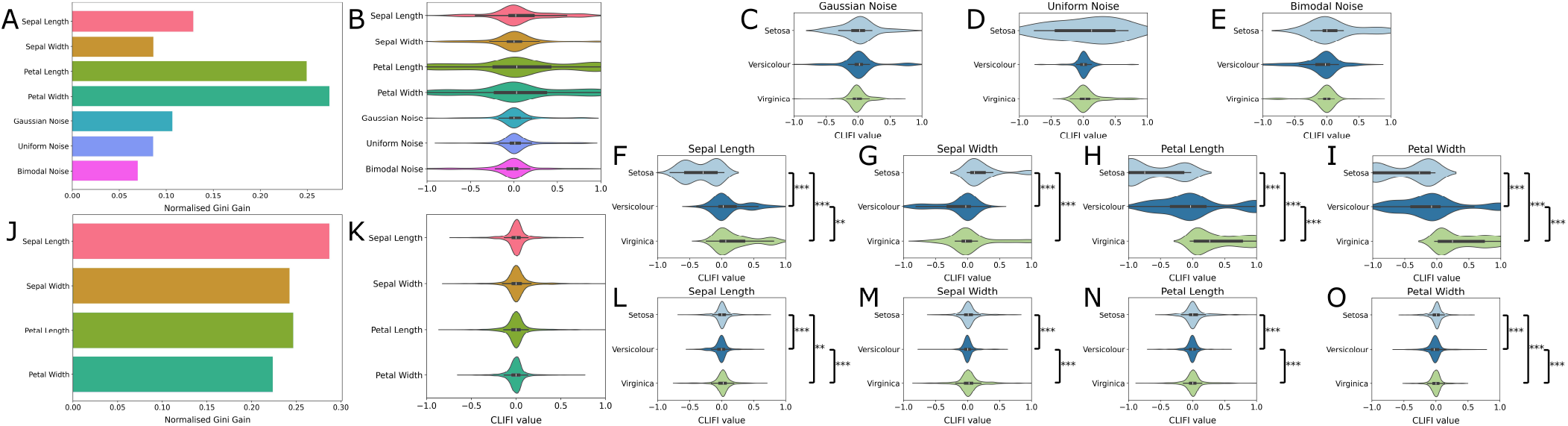
Visualisation of feature importance for Iris and Iris-permuted dataset for the RF model. (A) Normalised Gini coefficients for the Iris dataset. (B) Aggregated distribution of CLIFI values across all classes, CLIFI distributions that deviate from a normal distribution around 0 may have class information. (C) Distribution of CLIFI values for Gaussian noise. (D) Distribution of CLIFI values for uniform noise. (E) Distribution of CLIFI values for bimodal noise. (F) Distribution of CLIFI values for sepal length. (G) Distribution of CLIFI values for sepal width. (H) Distribution of CLIFI values for petal length. (I) Distribution of CLIFI values for petal width. (J) Normalised Gini coefficients for the Iris-permuted dataset. (K) Aggregated distribution of CLIFI values across all classes. (L) Distribution of CLIFI values for sepal length. (M) Distribution of CLIFI values for sepal width. (N) Distribution of CLIFI values for petal length. (O) Distribution of CLIFI values for petal width. Pairwise comparison within a class were tested. *** = p < 0.0005, ** = p < 0.005, * = p < 0.05.

While the Gini coefficients are still non-zero (Fig 1 J) for Iris-permuted, the CLIFI values are all centered around 0 (Fig 1K) indicating an equal split. The differences in the permuted features (Fig 1L-M) are all centered around 0 and considerably more narrow than those from the real model (Fig 1F-I), hence features with a wider dispersion of CLIFI values away from zero (toward either −1 or 1) signifies greater predictive importance in class separation.

#### TCGA Data - Feature Importance Visualisation

Analysis of the topmost important proteins across all cancers show that RF and LAVASET have the capability to detect proteins with similar average CLIFI magnitudes of importance across multiple cancer types, e.g. progesterone receptor (PR) in RF, and cyclin B1 and ER*α* (oestrogen receptor alpha) in both RF and LAVASET. We visualised the aggregated CLIFI values (normalised to the maximum value to allow comparison between methods) as heatmaps in Figure 2.

**Fig. 2.**
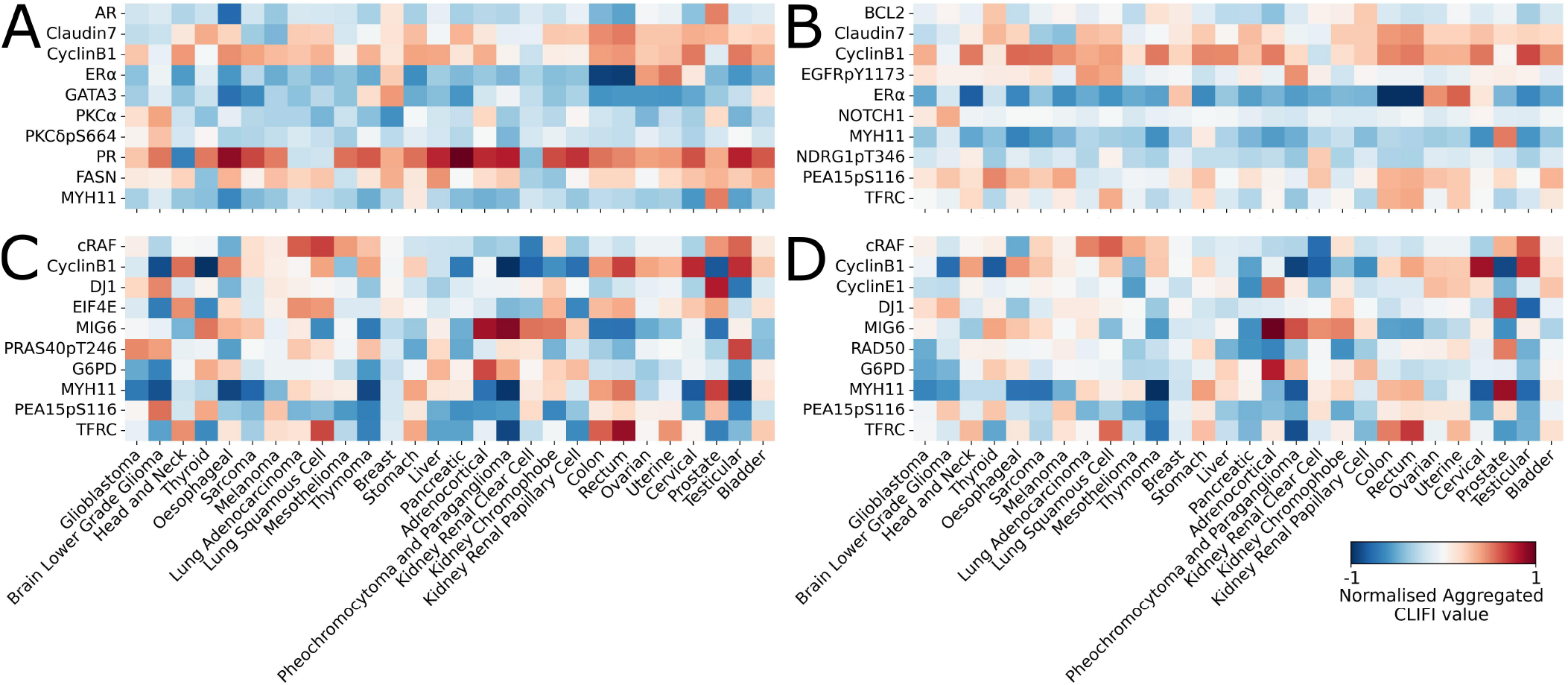
Heatmap of the top ten most important proteins across all 28 cancer types. Proteins selected based on the highest absolute normalised aggregated CLIFI value. (A) RF, (B) LAVASET, (C) GBDT, and (D) LAVABOOST. Colour bar shows the feature importance gradient, from negative to positive CLIFI-values, with white indicating a zero value.

Notably, within these proteins with high naCLIFI magnitude concordance, there are certain cancer types with significantly different naCLIFI values when compared to the majority. For example, Figure 2A shows that while the average naCLIFI values for PR is close to 1 in most cancer types, in lung adenocarcinoma, lung squamous cell, and kidney renal clear cell, the average naCLIFI value is close to −1. Additionally, Figure 2B shows that while the average naCLIFI value for Cyclin B1 is close to 1 in most cancer types, in kidney renal clear cell the average naCLIFI value is close to −1. However, this is not seen in the GBDT or LAVABOOST (Fig 2C,D), instead most proteins have a high CLIFI value magnitude for some cancer types while the others have varying magnitudes. For example, the MYH11 protein has a high magnitude of naCLIFI values for thymoma (<0) and prostate cancer (> 0) in all models, with the boosting models showing greater variability due to these predicting errors of prior trees.

#### TCGA Data - Interpretability of Feature Importance in Latent Variable Embedding Models

While the heatmaps show the overall importance, these plots fail to display the relations between features. LAVASET and LAVABOOST integrate topological information, hence the outputs are conditional on the mapping of features. Figure 3 shows the top proteins for four cancer types for which LAVASET outperformed all other methods. The edges show the neighbours in the distance matrix for calculating the latent embedding to facilitate interpreting the feature relations.

**Fig. 3.**
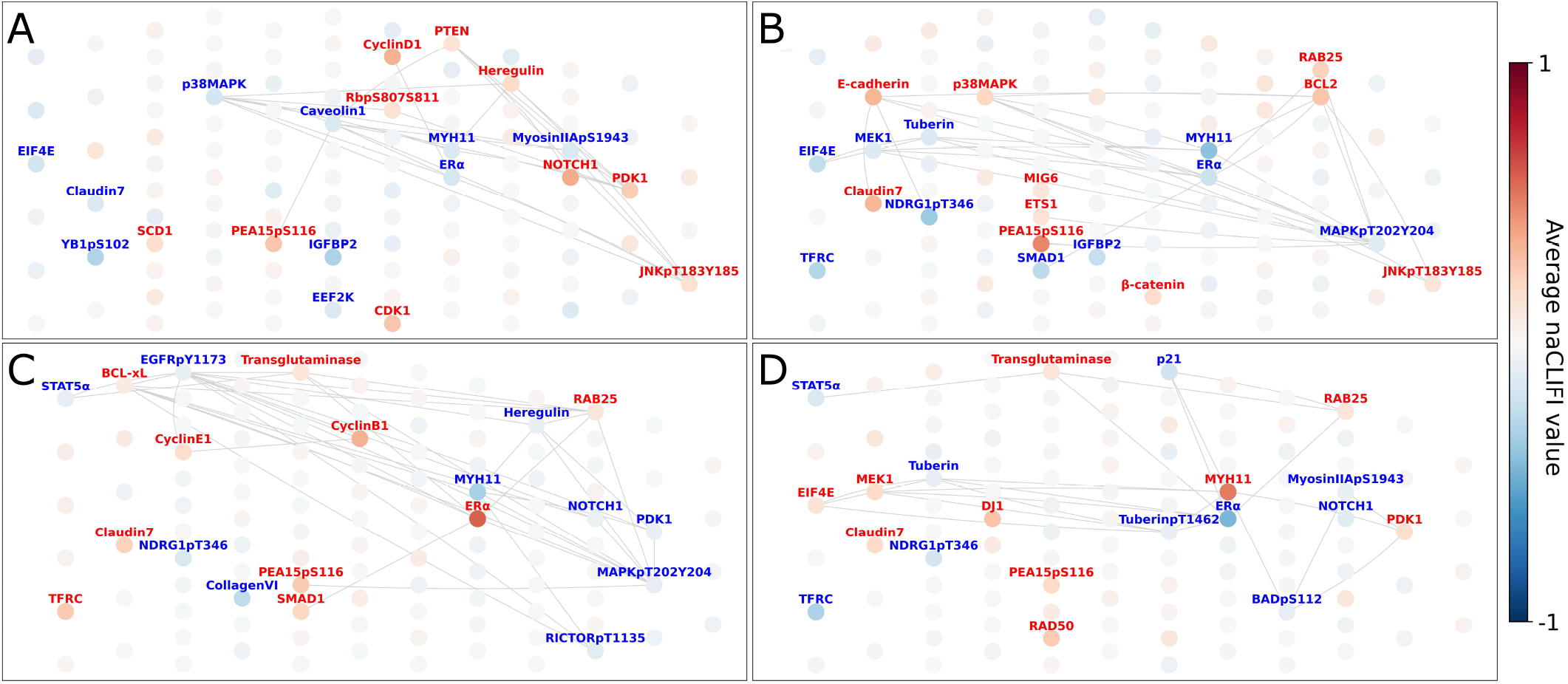
Visualisation of important features (node colour) and protein-protein interactions (edges) for selected cancer types from the LAVASET multi-class classification model. (A) Brain lower grade glioma (F1 = 0.97). (B) Thyroid cancer (F1 = 0.98). (C) Uterine endometrial cancer (F1 = 0.95). (D) Prostate adenocarcinoma (F1 = 1.00). Edges are only shown if labelled proteins have interactions.

It can be seen that while MYH11 (smooth muscle myosin heavy chain 11) has interactions with BCL2 (B-cell lymphoma 2) and p38MAPK (p38 mitogen-activated protein kinase), and ER*α* with RAB25 (Ras-related protein in brain 25) and SMAD1 (Mothers Against Decapentaplegic Homolog 1), this is independent from the signs of the naCLIFI values. Incorporating the topological information allows for interpretation of the model in terms of (joint) function and role rather than on an individual protein basis (as with RF and GBDT). More relations between proteins are incorporated into the model, however here we only visualise the top few for simplicity.

#### TCGA Data - Individual Feature Importance Evaluation

Similar to the Iris data, the CLIFI values allow for the visualisation of feature importance across all classes for individual features, with the distribution indicating the sign of association with the split (Fig 4). As example, we show several proteins with different distributions of CLIFI values.

**Fig. 4.**
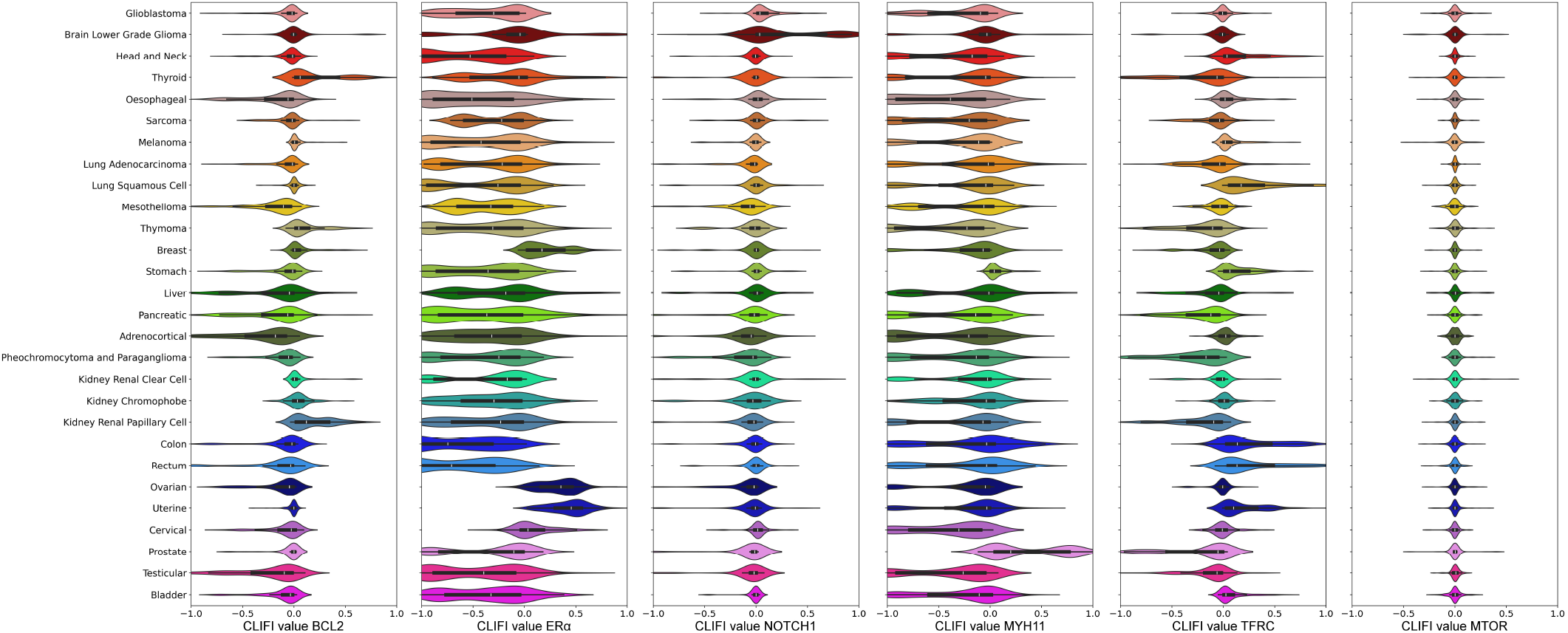
Visualisation with violin plots of five selected top proteins, and one unimportant, from the LAVASET multi-class classification model. From left to right: BCL2, ER*α*, NOTCH1, MYH11, TFRC, MTOR.

For example, the distribution of CLIFI values for BCL2 indicates that it is importance in the classification of several cancers, each of which appear to have bi- or multimodal distributions (hinting at heterogeneity), and specifically with thyroid and kidney renal papillary cell (higher expression) and oesophageal, mesothelioma, liver, pancreatic and adrenocortical (lower expression) cancers. TFRC (transferrin receptor) shows CLIFI distributions trending in opposite directions for lung squamous cell carcinoma (higher) and lung adenocarcinoma (lower) indicating potentially discriminatory potential between different cancers of the same organ.

Other proteins such as ER*α* and MYH11 have clear patterns for specific cancers with higher and lower expression of these proteins, with the bimodal distributions more evident that for other proteins. NOTCH1 (Neurogenic locus notch homolog protein 1) has CLIFI values largely centered around 0 except for brain lower grade gliomas, however while it may appear its expression may not relate to differences between other cancers, its distributions are considerably different from the MTOR (mammalian target of rapamycin) protein which is not associated with any class separation in any model.

We have observed that the CLIFI values relate to the protein expression levels in the original data. For example for MYH11 (Fig 5) prostate and stomach cancer have the highest CLIFI values matching the protein expression levels. Likewise, oesophageal, thymoma, cervical, and testicular cancers have CLIFI values indicating lower expression levels. Though mesothelioma, liver, prostate and some other cancer types exhibit bimodal (or multimodal) distributions pointing towards possible heterogeneity in the proteomic signature of these cancers.

**Fig. 5.**
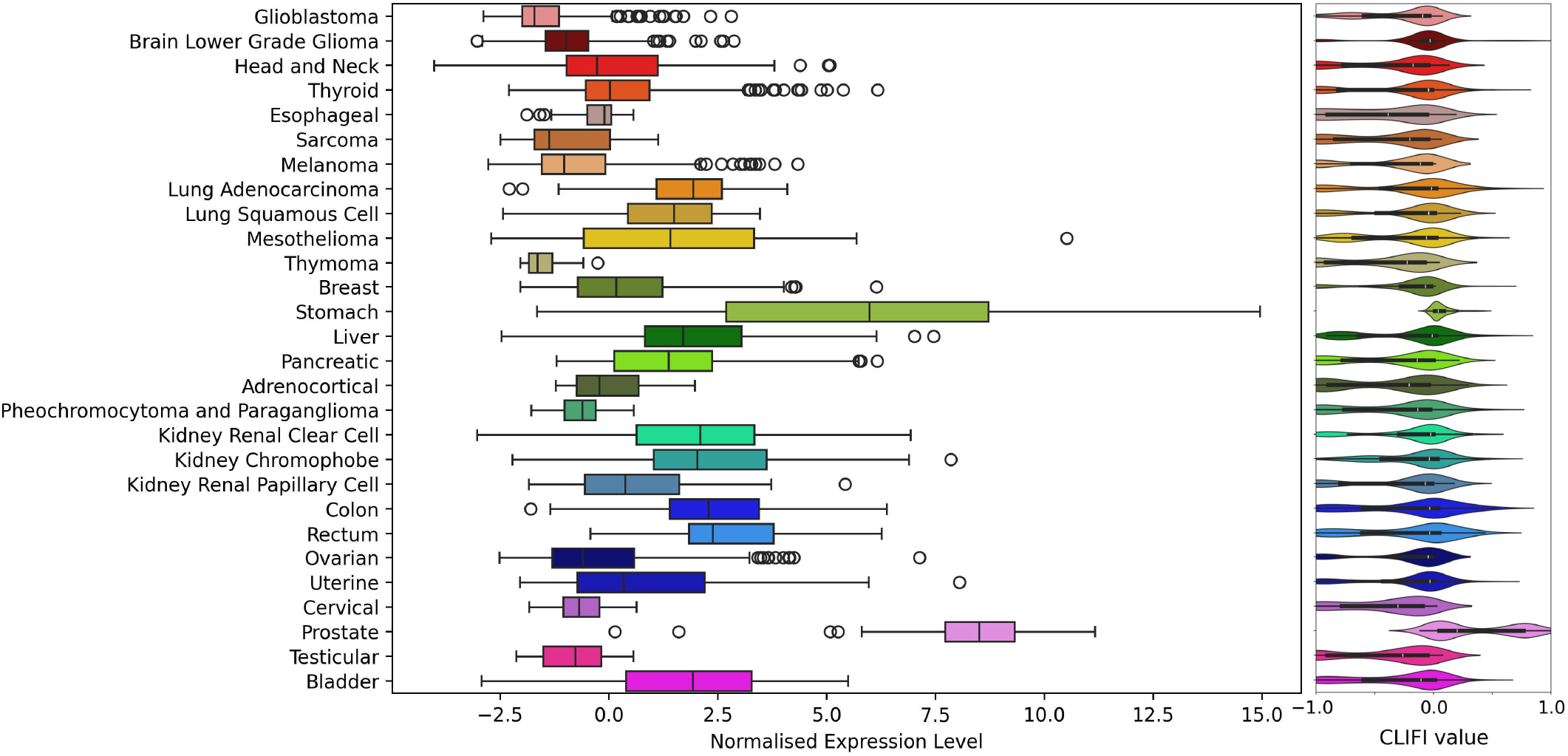
Comparison of protein expression level for MYH11 with CLIFI value from LAVASET model.

#### TCGA Data - Visualisation of Heterogenity in Classification of Cancer Types

The model performance is given with performance values for each cancer types, however this fails to show the homo- and heterogeneity of the data. We calculate a proximity matrix from the output from the ensemble model which represents the frequency that pairs of samples end up in the same leaf node in the classification model. We graphically represent the proximities as input to UMAP (2 dimensions) in Figure 6. The CLIFI values for several proteins indicated that breast cancer display bimodal distributions, and the UMAP plot shows that there appear to be multiple breast and stomach cancer subgroups based on the proteomics-based proximity matrix from LAVASET and visualises the similarity between cancer types based on the model predictions.

**Fig. 6.**
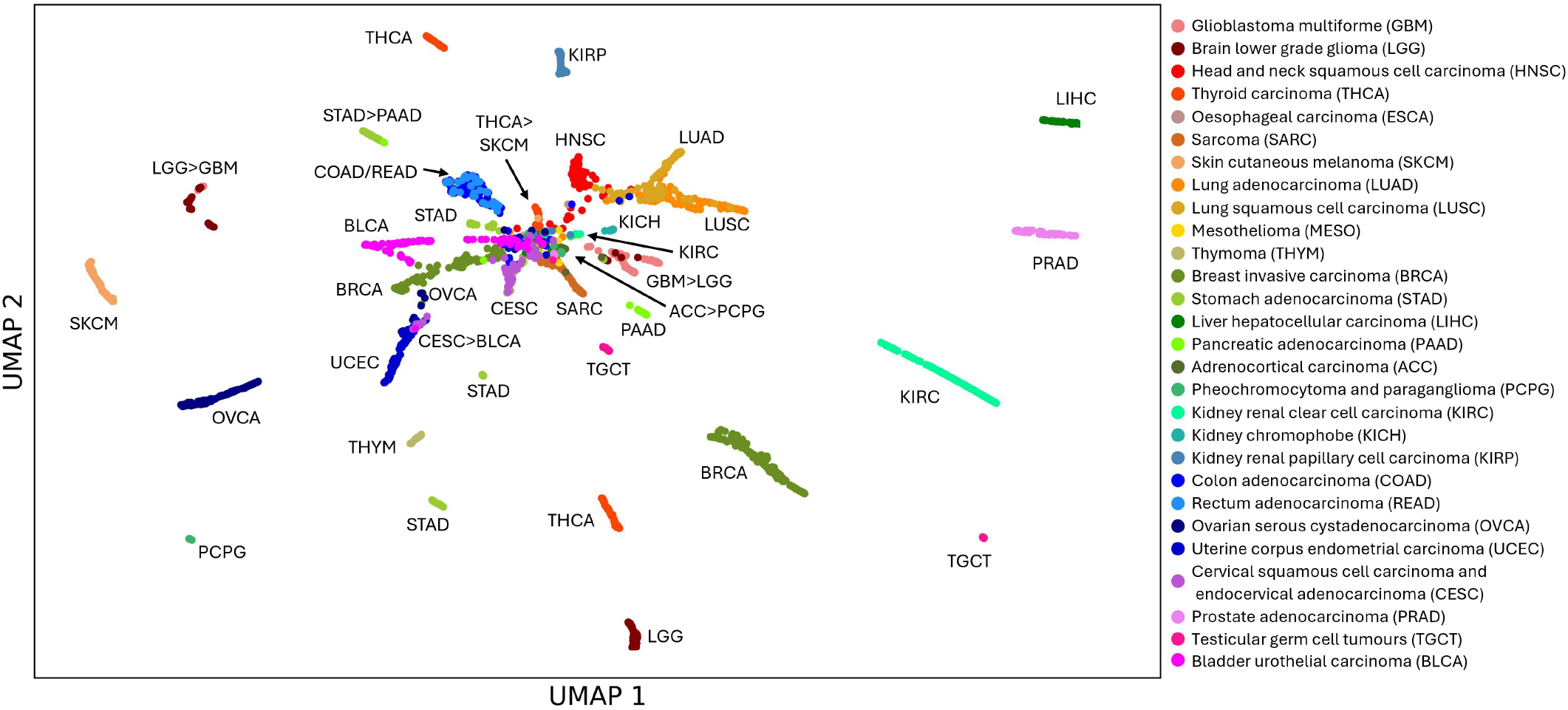
UMAP visualisation of the LAVASET proximity matrix with each cancer type labelled showing heterogeneity of some cancer types in the model prediction. *A* > *B* indicates cluster contains more samples from cancer type A than cancer B, ‘/’ indicates completely overlapped samples within a cluster.

While ovarian and uterine cancer had similar CLIFI values to breast cancer for some proteins (e.g. ER*α*), overall they cluster separately from breast cancer. The UMAP visualisation also shows that colon and rectal cancer (that are indistinguishable) are similar to two clusters of stomach cancer samples, the two lung cancers are closely related to head and neck cancers, and there are 2 clusters of brain cancers (one with majority of glioblastoma samples, another with majority of lower grade gliomas), all of which relates to the relative proximity of these cancers in the body.

## Discussion

We have introduced a new directional feature importance metric, CLIFI, that is integrated within the algorithm. The distribution of CLIFI values contain information on class separation and can be represented both on a global and class-specific level. Therefore, combined with being embedded within the training procedure, it provides unique insights that other methods such as the Gini coefficient and SHAP values do not, such as allowing the identification of heterogeneity within a class for specific features. Secondly, we have extended the LAVA framework to gradient boosting methods and introduced the LAVABOOST algorithm here for datasets with correlated features and multiple classes. While our results indicate that for these data RF and LAVASET outperform GBDTs and LAVABOOST, all methods at least outperform others for some cancer types in the TCGA proteomics data.

Since the CLIFI calculation is based on measuring the change in class distribution, it is not dependent on the class prediction accuracy and provides a new level of interpretability of stochastic ensemble models. Unlike the Gini coefficient, also arising from the training model, we demonstrated that CLIFI values are (significantly) higher for real features than random ones for the Iris example dataset, and that if the feature contains little class information (Iris-permuted) the CLIFI values have narrow intervals and center around 0, whereas Gini coefficients are still large. The same was shown for the real TCGA data where unimportant features have narrow CLIFI distributions around 0 (e.g. MTOR) and predictive features such as BCL2, ER*α*, NOTCH1, and MYH11 deviate from this. Although, this may suggest that CLIFI is correlated with accuracy, both RF and LAVASET assign a strikingly high importance to ER*α* for classifying rectum adenocarcinoma, with overall F1-scores of 0 and 4.65% for RF and LAVASET, respectively. CLIFI resembles the manner in which the model makes its predictions. In the case of rectum adenocarcinoma, these patients are predicted with colon adenocarcinoma patients since both have very similar importance assignments in the heatmaps and the precision of colon adenocarcinoma is noticeably lower compared to other cancers (71-72% versus 84-100%). Another benefit of using a feature importance calculation based on the change of class distribution is that it is unaffected by unbalanced classes, unlike other commonly used methods such as the Gini coefficient and SHAP values. Finally, the CLIFI distribution identifies the feature profile used by the model for specific class predictions, and it can be used to identify features with bi- or multimodal distributions within a class, even though overall the class could appear as homogenous.

While our aim here was to demonstrate the new methodology, the use of the publicly available TCGA data (26) allow comparison of model outputs with previously published literature. For example, for uterine corpus endometrial carcinoma LAVASET (25) outperforms the other methods and the proteins with the highest magnitude of average naCLIFI values (MYH11, ER*α*, and cyclin B1) have all been found to be expressed in the same direction that is indicated by the CLIFI values by others in endometrial cancers (31–33) previously.

The original LAVASET method was applied on different datasets, but with distances defined by the data itself (spatial or temporal) (25). Here we integrated protein interaction data into this step to allow the method to make the decisions based on topological (pathway) information. Analysis of the top proteins with the most interactions identifies pathways which are also important for disease progression in cancer, such as epidermal growth factor signalling (via EGFR and EGFRpY1173) (34), the MAPK pathway (via p38MAPK and MAPKpT202Y204) (35), and BCL-regulated apoptotic pathway (via BCL-2 and BCL-xL) (36). However, not all proteins associated with each pathway are identified a s being important. Since we used a multi-class model, the feature importance profile will highlight the most differentiating proteins, and thus will not be an exact match to proteomic profiles identified in biological studies. Additionally, proteins can be part of multiple pathways, hence this impacts on the ‘neighbours’ used to calculate the latent feature embedding in ‘LAVA’.

While the UMAP visualisation of model proximities has shown some heterogeneity and subclasses within some cancer types (e.g. breast and stomach cancer), analysis of the CLIFI values allows to identity the features responsible for this (bimodal distributions of ER*α* and others). However, despite MYH11 being the most important protein for prostate cancer prediction and the CLIFI values indicating a bimodal distribution, the UMAP shows only one cluster. While differential ER*α* expression has been reported in prostate cancer patients (37) however, to the best of our knowledge, differential MYH11 expression has not thereby warranting further investigation. The CLIFI distribution will allow further a posteriori analysis, such as tests for normality, deviation from 0, and others using conventional, univariate statistical tests. When comparing CLIFI values between models as done here, they should be normalised. Whereas for the more common case where only a single model is calculated, the aggregated CLIFI values can be used on their own.

We have also extended the ‘LAVA’ framework to boosting methods, but observed that LAVASET outperformed LAVABOOST, therefore. However, for these data, RF and LAVASET appear to be the most predictive models, hence their CLIFI values are potentially of more interest. However, CLIFI can be implemented in other methods such as XGBoost (38) which may have higher performance. For our data, RF and LAVASET had the highest performance with F1-scores of 92.29% and 91.83%, respectively. However, optimising the additional parameters of XGBoost (sample random state) had an F1 of 92.53%. Since the LAVABOOST outperformed GBDT, it is plausible that the performance of XGBoost may increase further with the addition of the ‘LAVA’ step and the integration of CLIFI will improve its interpretability.

In summary, we have contributed an integrated, directional feature importance metric for decision tree-based models to facilitate feature importance assessment for multi-class classification. This metric can be used together with incorporating topological information into the decision functions of tree-based algorithms. The incorporation of topological information has been extended here to protein interactions but can be used with any type of network information that links together individual features to add inductive bias into the model for improved interpretability. Finally, several findings from analysing the individual directional feature importances match with existing literature, while other observations have not, generating potentially new testable hypotheses for cancer heterogeneity and subsequent clinical diagnostic studies.

## AUTHOR CONTRIBUTIONS

Conceptualisation, M.K. and J.M.P.; Data curation, E.R.L. and S.M.Y.; Formal analysis, E.R.L. and S.M.Y.; Funding acquisition, J.M.P.; Investigation, E.R.L. and S.M.Y.; Methodology, M.K. and J.M.P.; Project administration, M.K. and J.M.P.; Resources, J.M.P.; Software, E.R.L. and M.K.; Supervision, M.K. and J.M.P.; Validation, E.R.L. and J.M.P.; Visualisation, E.R.L. and J.M.P.; Writing – original draft, E.R.L.; Writing – review & editing, M.K. and J.M.P. All authors have read and agreed to the published version of the manuscript.

## Appendix Class-specific Classification Performance

**Table 3.** Classification report of RF and LAVASET when classifying 28 cancer types for a single initialisation. Values in bold indicate the highest performance (expressed as percentage) for each metric per cancer type across the 4 methods (see Table 4 for results for GBDT and LAVABOOST).

| Cancer type | Precision |  | Recall |  | F1-score |  | Support |
| --- | --- | --- | --- | --- | --- | --- | --- |
|  | RF | LAVASET | RF | LAVASET | RF | LAVASET |  |
| Glioblastoma | 95.77 | 95.71 | **93.15** | 91.78 | **94.44** | 93.71 | 73 |
| Brain lower grade glioma | **96.97** | **96.97** | 97.71 | 97.71 | **97.34** | **97.34** | 131 |
| Head and Neck | **94.59** | 89.66 | **98.13** | 97.2 | **96.33** | 93.27 | 107 |
| Thyroid | **96.64** | **96.64** | **100** | **100** | **98.29** | **98.29** | 115 |
| Esophageal | **77.27** | 75 | **89.47** | 86.84 | **82.93** | 80.49 | 38 |
| Sarcoma | **90.67** | 85.71 | **100** | 97.06 | **95.1** | 91.03 | 68 |
| Melanoma | 95.28 | 98 | **95.28** | 92.45 | **95.28** | 95.15 | 106 |
| Lung adenocarcinoma | 84.96 | **85.71** | 87.27 | 87.27 | 86.1 | **86.49** | 110 |
| Lung squamous cell | **85.57** | 83.67 | **83.84** | 82.83 | **84.69** | 83.25 | 99 |
| Mesothelioma | **100** | **100** | **73.68** | **73.68** | **84.85** | **84.85** | 19 |
| Thymoma | **100** | **100** | 92.59 | **96.3** | 96.15 | **98.11** | 27 |
| Breast | **95.45** | 95.12 | **98.91** | **98.91** | **97.15** | 96.98 | 276 |
| Stomach | 94.64 | 95.5 | **98.15** | **98.15** | 96.36 | 96.8 | 108 |
| Liver | **92.86** | 89.09 | **92.86** | 87.5 | **92.86** | 88.29 | 56 |
| Pancreatic | **96.77** | 90.91 | **83.33** | **83.33** | **89.55** | 86.96 | 36 |
| Adrenocortical | 92.31 | **100** | **85.71** | **85.71** | 88.89 | **92.31** | 14 |
| Pheochromocytoma and Paraganglioma | 95.83 | **100** | 92 | 96 | 93.88 | 97.96 | 25 |
| Kidney renal clear cell | **98.61** | 96.6 | **98.61** | **98.61** | **98.61** | 97.59 | 144 |
| Kidney chromophobe | **100** | 94.44 | **94.74** | 89.47 | **97.3** | 91.89 | 19 |
| Kidney renal papillary cell | 90.91 | 92.42 | 92.31 | **93.85** | 91.6 | **93.13** | 65 |
| Colon | 71.53 | **72.14** | **94.5** | 92.66 | **81.42** | 81.12 | 109 |
| Rectum | 0 | **33.33** | 0 | **2.5** | 0 | **4.65** | 40 |
| Ovarian | **99.21** | 98.4 | **96.92** | 94.62 | **98.05** | 96.47 | 130 |
| Uterine endometrial | **95.31** | 94.03 | 92.42 | **95.45** | 93.85 | **94.74** | 132 |
| Cervical | **97.37** | 97.3 | **71.15** | 69.23 | **82.22** | 80.9 | 52 |
| Prostate | **100** | **100** | **100** | **100** | **100** | **100** | 106 |
| Testicular germ cell | 94.87 | 97.3 | **100** | 97.3 | 97.37 | 97.3 | 37 |
| Bladder urothelial | 93.27 | 95.1 | **94.17** | **94.17** | 93.72 | **94.63** | 103 |
| Weighted accuracy | 91.92 | **91.97** | **93.01** | 92.49 | **92.29** | 91.83 | 2345 |

**Table 4.**
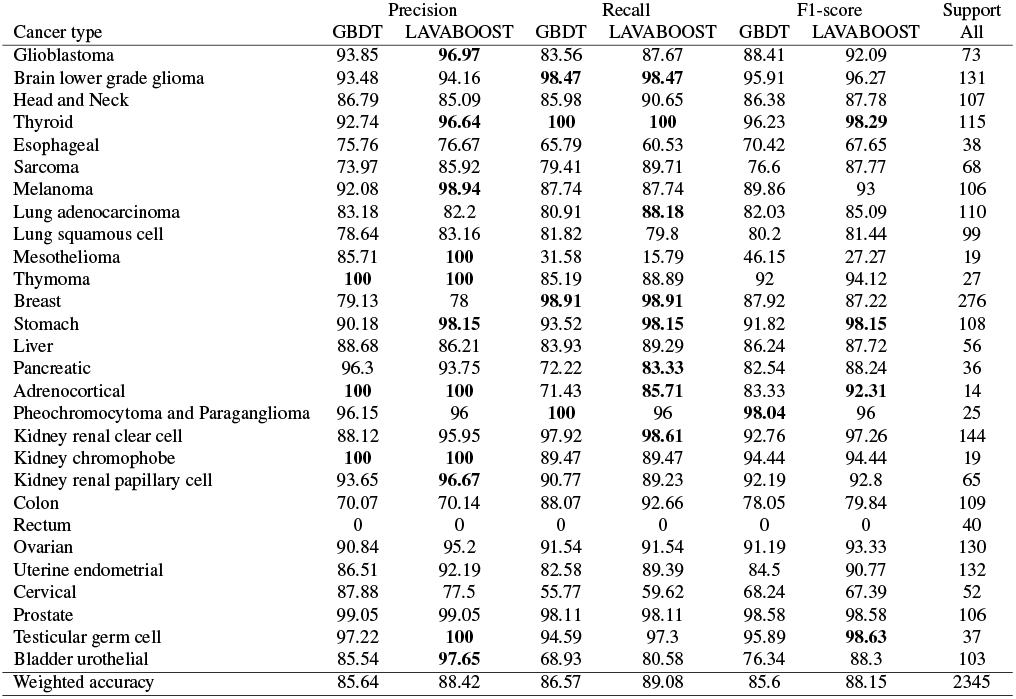
Classification report of GBDT and LAVABOOST when classifying 28 cancer types for a single initialisation. Values in bold indicate the highest performance (expressed as percentage) for each metric per cancer type across the 4 methods (see Table 3 for results for RF and LAVASET).

| Cancer type | Precision |  | Recall |  | F1-score |  | Support |
| --- | --- | --- | --- | --- | --- | --- | --- |
|  | GBDT | LAVABOOST | GBDT | LAVABOOST | GBDT | LAVABOOST |  |
| Glioblastoma | 93.85 | **96.97** | 83.56 | 87.67 | 88.41 | 92.09 | 73 |
| Brain lower grade glioma | 93.48 | 94.16 | **98.47** | **98.47** | 95.91 | 96.27 | 131 |
| Head and Neck | 86.79 | 85.09 | 85.98 | 90.65 | 86.38 | 87.78 | 107 |
| Thyroid | 92.74 | **96.64** | **100** | **100** | 96.23 | **98.29** | 115 |
| Esophageal | 75.76 | 76.67 | 65.79 | 60.53 | 70.42 | 67.65 | 38 |
| Sarcoma | 73.97 | 85.92 | 79.41 | 89.71 | 76.6 | 87.77 | 68 |
| Melanoma | 92.08 | **98.94** | 87.74 | 87.74 | 89.86 | 93 | 106 |
| Lung adenocarcinoma | 83.18 | 82.2 | 80.91 | **88.18** | 82.03 | 85.09 | 110 |
| Lung squamous cell | 78.64 | 83.16 | 81.82 | 79.8 | 80.2 | 81.44 | 99 |
| Mesothelioma | 85.71 | **100** | 31.58 | 15.79 | 46.15 | 27.27 | 19 |
| Thymoma | **100** | **100** | 85.19 | 88.89 | 92 | 94.12 | 27 |
| Breast | 79.13 | 78 | **98.91** | **98.91** | 87.92 | 87.22 | 276 |
| Stomach | 90.18 | **98.15** | 93.52 | **98.15** | 91.82 | **98.15** | 108 |
| Liver | 88.68 | 86.21 | 83.93 | 89.29 | 86.24 | 87.72 | 56 |
| Pancreatic | 96.3 | 93.75 | 72.22 | **83.33** | 82.54 | 88.24 | 36 |
| Adrenocortical | **100** | **100** | 71.43 | **85.71** | 83.33 | **92.31** | 14 |
| Pheochromocytoma and Paraganglioma | 96.15 | 96 | **100** | 96 | **98.04** | 96 | 25 |
| Kidney renal clear cell | 88.12 | 95.95 | 97.92 | **98.61** | 92.76 | 97.26 | 144 |
| Kidney chromophobe | **100** | **100** | 89.47 | 89.47 | 94.44 | 94.44 | 19 |
| Kidney renal papillary cell | 93.65 | **96.67** | 90.77 | 89.23 | 92.19 | 92.8 | 65 |
| Colon | 70.07 | 70.14 | 88.07 | 92.66 | 78.05 | 79.84 | 109 |
| Rectum | 0 | 0 | 0 | 0 | 0 | 0 | 40 |
| Ovarian | 90.84 | 95.2 | 91.54 | 91.54 | 91.19 | 93.33 | 130 |
| Uterine endometrial | 86.51 | 92.19 | 82.58 | 89.39 | 84.5 | 90.77 | 132 |
| Cervical | 87.88 | 77.5 | 55.77 | 59.62 | 68.24 | 67.39 | 52 |
| Prostate | 99.05 | 99.05 | 98.11 | 98.11 | 98.58 | 98.58 | 106 |
| Testicular germ cell | 97.22 | **100** | 94.59 | 97.3 | 95.89 | **98.63** | 37 |
| Bladder urothelial | 85.54 | **97.65** | 68.93 | 80.58 | 76.34 | 88.3 | 103 |
| Weighted accuracy | 85.64 | 88.42 | 86.57 | 89.08 | 85.6 | 88.15 | 2345 |

